# A mathematical model of the disruption of glucose homeostasis in cancer patients

**DOI:** 10.1101/2023.03.15.532725

**Authors:** Noah Salentine, Jonathan Doria, Chinh Nguyen, Gabriella Pinter, Shizhen Emily Wang, Peter Hinow

## Abstract

In this paper we investigate the disruption of the glucose homeostasis at the whole-body level by the presence of cancer disease. Of particular interest are the potentially different responses of patients with or without hyperglycemia (including Diabetes Mellitus) to the cancer challenge, and how tumor growth, in turn, responds to hyperglycemia and its medical management. We propose a mathematical model that describes the competition between cancer cells and glucose-dependent healthy cells for a shared glucose resource. We also include the metabolic reprogramming of healthy cells by cancer-cell-initiated mechanism to reflect the interplay between the two cell populations. We parametrize this model and carry out numerical simulations of various scenarios, with growth of tumor mass and loss of healthy body mass as endpoints. We report sets of cancer characteristics that show plausible disease histories. We investigate parameters that change cancer cells’ aggressiveness, and we exhibit differing responses in diabetic and non-diabetic, in the absence or presence of glycemic control. Our model predictions are in line with observations of weight loss in cancer patients and the increased growth (or earlier onset) of tumor in diabetic individuals. The model will also aid future studies on countermeasures such as the reduction of circulating glucose in cancer patients.

## 1 Introduction

In recent years it has become increasingly clear that cancer is associated with a wide set of risk factors and co-morbidities that aggravate the clinical outcomes, undermine the benefits of cancer therapy, and contribute to disparities in cancer mortality [1]. Prominent among these is an impaired glucose/insulin homeostasis that manifests itself in insulin resistance (IR). Rapidly proliferating cancer cells require large amounts of energy and building materials. A large portion of these come in form of glucose, while other nutrients such as amino acids and lipids also contribute to cancer cell energetics and proliferation. On one hand, cancer cells enhance the rate of nutrient consumption and exhibit an altered metabolic pattern to meet their demands for biosynthesis as a new and fast-growing organ in the body. On the other hand, cancer cells also reprogram the metabolism of their host organism to further tip the balance of energy in their favor [2, 3]. There is currently an intensive research effort into the precise mechanisms by which the cancer cells achieve this. For example it has been shown that extracellular vesicles secreted by breast cancer cells and carrying miRNA suppress glucose consumption by brain and lung cells [4] and also impair insulin secretion to exert a broad effect on insulin-responsive tissues [5].

In healthy individuals, blood glucose levels are tightly regulated to achieve homeostasis. A disruption of this regulation mechanism may result in conditions such as Diabetes Mellitus. The glucose-insulin-glucagon feedback system has been studied by physiologists, mathematicians and biomedical engineers for nearly 60 years. This has resulted in an impressive number of mathematical models and experimental studies, see [6] for a review. In order to keep the complexity at a manageable level, in this work we forgo the explicit modeling of insulin and the precise mechanism by which some cancer may desensitize insulin response in the healthy peripheral cells. Instead we focus on the interplay between different cell populations competing for glucose.

Here we propose an ordinary differential equation (ODE) model for the competition between a cancer cell population and the healthy glucose-dependent remaining body. The conceptual model is depicted in Figure 1. Its central part is a denial of glucose to the healthy part of the body in the presence of cancer. This is a combined consequence of the simultaneous consumption of glucose by the two parties from a common glucose reservoir, and the ability of some cancer cells to suppress glucose consumption by healthy tissues. We investigate the model through numerical simulations and exploring the parameter space. Our results are of qualitative nature. For example, what is the influence of parameters on the time to the tumor reaching a certain size and on body mass loss associated with glucose reallocation?

**Fig. 1.**
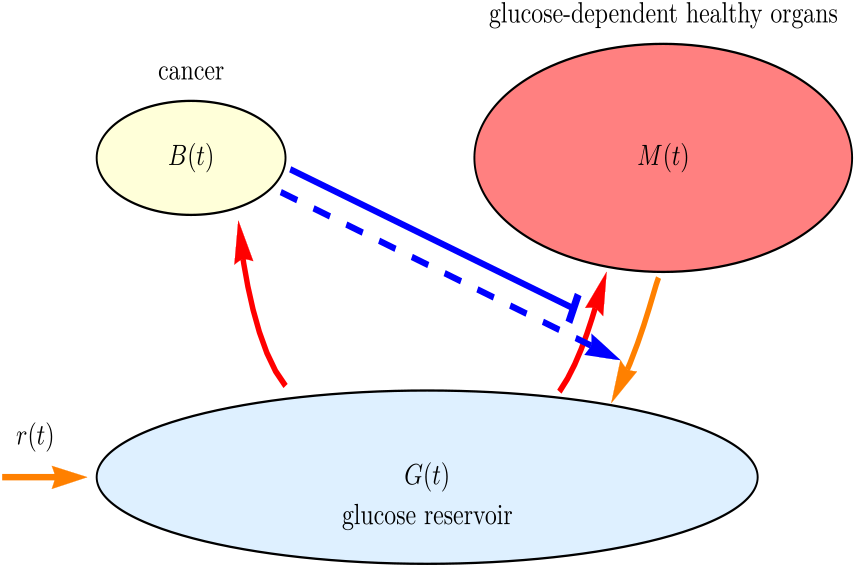
The competing cell populations (top row) and their common glucose resource. In this model, some cancer cells could suppress the glucose consumption by the glucose-dependent healthy body mass.

Moreover, we investigate how the disease differs in diabetic respectively nondiabetic patients. Another aspect we address is the possible use of drugs such as Metformin as supportive therapy for cancer patients [7–9].

## 2 The mathematical model

We begin by considering a cancer-free human in whom glucose uptake, loss and body mass are in a stable equilibrium. The variables of our model are

1. *G*(*t*), the amount of available glucose to support the body at time *t*, (in g)
2. *M*(*t*), the glucose-dependent body mass of the individual (in g), and
3. *B*(*t*), the mass of the cancer tissue (in g), which will be included in the second step of the model construction.

The free, accessible form of circulating glucose exists in a reservoir from which the whole body draws and which is replenished regularly by consumption of food. There are various ways this reservoir can be defined, depending for example whether glycogen stored in the liver and skeletal muscle is counted towards it. For the sake of simplicity, we only count the amount of glucose constantly circulating in the blood. For a non-diabetic person this is approximately 4 g [10], while it can be up to 11 g for a diabetic person [11]. At the tissue or organ level, some tissues, such as muscle and adipose tissue, exhibit a high degree of plasticity as their size and weight fluctuate dramatically in response to the availability of nutrients such as glucose. In contrast, other tissues such as bone and brain exhibit a lower degree of fluctuation in size and weight in response to nutrient conditions. Here we focus on those tissues that are highly responsive to glucose availability and their contribution to the overall body mass, which we refer to “glucose-dependent body mass” in this paper.

Glucose is supplied through food, consumed by the body, and lost at a certain rate through excretion by the kidneys. As our first equation, we propose

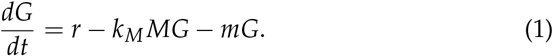

On a time scale of days, the supply rate *r* is time-dependent, and roughly 24 h-periodic in response to meals and circadian rhythm. However, on a timescale of months or years, as usually used in the clinical follow-up of cancer patients, we ignore the daily spikes and assume a constant supply rate *r*. Glucose is lost at a rate *m*. The constant models the consumption of the glucose by the body (the subscript *M* is used to distinguish it from a similar constant for the cancer to be introduced later). It is mirrored by a corresponding term in the equation for the glucose-dependent body mass

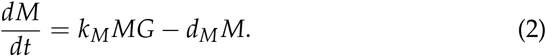

The constant *d_M_* models the rate of glucose consumption due to the regular metabolic processes of the body that are independent of building up body mass. In Equation (2) it takes the form of a mass loss rate for the case that no glucose is resupplied. We assume this is the body mass minus the mass of the skeleton.

We find the equilibrium solution of the model (1)-(2) to be

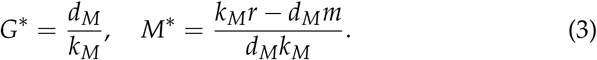

Since *M*\* > 0, this requires that the parameters satisfy the inequality *k_M_r* > *d_M_m*. This equilibrium can be shown to be asymptotically stable. That is, a small perturbation away from it will disappear in time.

When cancer is present, we modify the glucose balance Equation (1) to include two new phenomena. Firstly, we add a term to model the glucose consumption by the cancer cells. For this we use a term similar to — *k_M_MG* in Equation (1), and just replace *M* by *B* and introduce another constant. Secondly, we assume that the glucose consumption rate by the healthy tissue decreases in the presence of the cancer, such as through the mechanisms previously reported [4, 5]. Thus we have to replace Equation (1) by

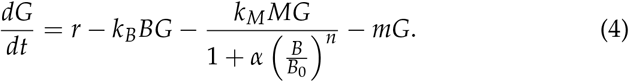

Here *k_B_* is the glucose consumption rate of the cancer. Once the cancer reaches a critical mass *B*_0_, it begins to impair the glucose consumption. The strength of this suppression is encoded in the constants *α* and *n*. Equation (2) is altered accordingly so that it becomes

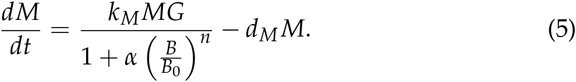

The cancer cells grow depending on the availability of glucose and die at a rate *d_B_* > 0. Thus

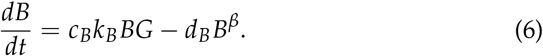

The conversion factor *c_B_* allows for the fact that the cancer uses glucose differently than the healthy cells with regard to growing its mass. The exponent *β* accounts for an increased mortality at larger cell numbers. This could be caused for example by lack of oxygen and nutrients at the core of the cancer due to insufficient blood supply.

## 3 Implementing the model

### 3.1 Parameter choices in the cancer-free case

The following section is devoted to finding parameters first for the cancer free case. A guiding principle are the steady state values for *G*\* and *M*\*. A difficulty is that the value of *M*\* is not the same in every healthy subject as there is a wide variation of body weights. We choose *M*\* = 6 · 10^4^ g and, as stated before, *G*\* = 4 g in a non-diabetic person [10].

To determine the rate of glucose intake we consider a diet of 10,000 kJ per day for a healthy person. According to Institute of Medicine (IOM), the recommended daily allowance for carbohydrates is 130 g [12]. This is based on the necessity of glucose acting as the required fuel for the central nervous system [13, Section 30.2]. Additionally, it is recommended by the IOM that 45-65 % of the total energy intake is in form of carbohydrates. Using the chosen diet above, we have a range of 139 - 202 g of carbohydrates daily. We choose the center point of this interval, *r* = 164 g d^−1^ (±38.1 g d^−1^). The National Institutes of Health (NIH) and the American Diabetes Association (ADA) recommend low carbohydrate eating plans for individuals with type 2 diabetes [14, 15]. We implement this through a value of *r* that is reduced by 85 %, or to 140 g d^-1^ in diabetic patients. We also assume that all carbohydrates acquired from diet, regardless of their form, break down into glucose with a 100 % efficiency.

In healthy individuals an amount of up to 25 mg/dL glucose in urine is considered normal whereas higher values indicate glycosuria [16]. The range of urine volume is 800 to 2000 mL d^−1^. Thus about 0.2 - 0.5 g of the 4 g circulating glucose are lost daily. The center point of this interval is *m* = 0.0875 d^−1^ (±0.0375 d^−1^).

The glucose consumption rate is not easily available in the literature. We use the fact that at equilibrium,

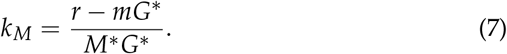

Using the values that we have already selected, we find that *k_M_* = 6.8 · 10^−4^ g^−4^d^−1^ for a non-diabetic person and *k_M_* = 2.1 · 10^−4^ g^−4^d^−1^ for a diabetic person.

The rate of mass loss *d_M_* can be estimated from various reports of the effects of starvation and semi-starvation experiments. [17] reported a 25 % loss of body weight in a six-month semi-starvation study on human volunteers, although the actual degree to which energy intake was reduced is not known. We work with the values given in Table 1 selected to match the equilibrium condition from Equation (3),

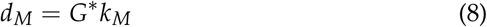

in cancer-free individuals.

**Table 1.** Parameters of the cancer-free model (1)-(2) in non-diabetic (column A) and diabetic (column B) individuals.

| parameter | value A | value B | role |
| --- | --- | --- | --- |
| $r$ | 164 g d<sup>-1</sup> | 140 g d<sup>-1</sup> | rate of glucose replenishment |
| $k_M$ | $6.8 \cdot 10^{-4} \text{ (g d)}^{-1}$ | $2.1 \cdot 10^{-4} \text{ (g d)}^{-1}$ | glucose consumption rate |
| $m$ | 0.0875 d<sup>-1</sup> | same | rate of glucose loss |
| $d_M$ | $2.7 \cdot 10^{-3} \text{ d}^{-1}$ | $2.3 \cdot 10^{-3}$ | rate of body mass loss |

### 3.2 Introduction of the cancer

Our choice of the parameters in a cancer patient is given in Table 2. As they are not available in the literature, their overall justification is that we are able to replicate plausible developmental histories of cancer. However, some comments can still be made. For example, we choose as a baseline value that cancer cells consume glucose at twice the rate of healthy cells in a non-diabetic individual. Th e higher glucose consumption rate in cancer cells is achieved partially through cancer-specific high expression of transporters (e.g., certain types of glucose transporters) and enzyme isoforms (e.g., hexokinase 2) that drive glucose flux forwards [18,19]. Glucose uptake in cancer and normal cells has been evaluated and compared by PET and 2-deoxyglucose incorporation, indicating 2-3 fold higher uptake in cancer cells that correlates with the expression levels of glucose transporters and hexokinase [20]. We choose

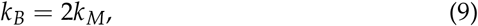

but this will be subjected to changes later on. Another constant that deserves some discussion is the dimensionless constant *c_B_* which is the way in which the cancer cells convert glucose into their maintenance and growth relative to the healthy cells. It is well known that they are inefficient when it comes to converting glucose into energy in form of ATP due to the Warburg effect [21]. However this does not necessarily imply that *c_B_* < 1 as the biosynthesis of proteins can actually be more efficient than in healthy cells. We will explore this deeper by varying this parameter below.

**Table 2.** Baseline parameters in the presence of cancer. Note that some parameters are varied in subsequent simulations.

| parameter | value | role |
| --- | --- | --- |
| $k_B$ | $1.4 \cdot 10^{-3} \text{ (g d)}^{-1}$ | rate of glucose consumption |
| $B_0$ | 10 g | cancer threshold size for glucose deprivation |
| $n$ | 0.3 | strength of the glucose suppression |
| $c_B$ | 0.35 | efficiency of glucose use by the cancer |
| $d_B$ | $1.2 \cdot 10^{-3} \text{ d}^{-1}$ | death rate of cancer cells |
| $\alpha$ | 1 | modulation of cancer competition for glucose |
| $\beta$ | 1.1 | exponent of cancer loss term |

With these parameters, we perform numerical simulations of the Equations (4)–(6). The initial condition for the cancer mass is *B*(0) = 0.5 g, while for *G*(0) and *M*(0) we use their equilibrium values in a diabetic respectively non-diabetic individual (which are the same for *M*(0)). In Figure 2 we show simulation results for a pair of non-diabetic patients. For 6-7 years there is no visible change until the body mass begins to drop and the cancer mass grows dramatically. We also observe an increase of the glucose circulating in the blood. As the glucose supply remains constant and the cancer suppresses the uptake of glucose by the healthy cells, the cancer does not react with increased consumption to the greater availability of glucose. Decreasing the parameter *α* below 1 results in a delay of the cancer eruption, which can be identified with a less aggressive cancer type. In Figure 3 we show simulation results for a non-diabetic and a diabetic patient. Clearly, the eruption of cancer is much earlier in the diabetic patient. We define the time *t*^*^ to be the time at which *B*(*t*\*) = 100 g. This choice of cancer mass to define the critical time point is somewhat arbitrary, as is the choice of the initial mass. However, at the qualitative level, the dependence of the time a threshold on the parameters is independent of the choice of the threshold. We report the doubling time at *t*\*,

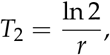

where *r* is the growth rate obtained by fitting the logarithm of the cancer mass *B*(*t*) up to *t*\* to a linear function. To quantify the mass loss, we select *M*(0) - *M*(*t*\*) and the rate -*M*^1^ (*t*\*) at that time. Finally, we report the elevation of blood glucose levels over their baseline values. These simulated cancer characteristics for the different simulation scenarios are collected in Table 3.

**Fig. 2.**
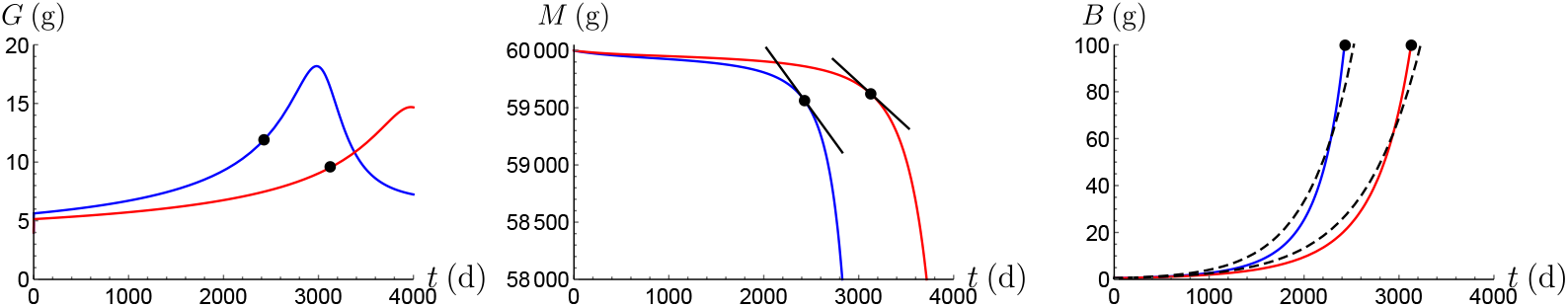
The amount of circulating glucose *G* (left), the glucose-dependent body mass *M* (center) and the cancer mass *B* (right) over a period of approximately 12 years. Blue lines indicate *α* = 1 (scenario “A”, a cancer more aggressively competing for glucose) and red lines indicate *α* = 0.7 (scenario “B”, a cancer less aggressively competing for glucose). In the first case we have that *t*\* = 2427 d, in the second case *t*\* = 3128 d. These time points are marked by black dots on the curves. The dashed lines in the right panel show the optimal fit with an exponential function which is used to determine the overall doubling time. Note that *M* is plotted on a reduced scale for better visibility.

**Table 3.** Cancer characteristics for the three simulated scenarios in Figures 2 and 3. From left to right are 1. the time for the cancer mass *B*(*t*) to reach 100 g, 2. the overall doubling time, 3. the total mass loss, 4. the rate of mass loss at the time *t*\*, and 5. the elevation of the blood glucose level compared to the normal level.

| scenario | $t^*$ (d) | $T_2$ (d) | $M(0) - M(t^*)$ (g) | $-M'(t^*)$ (g d<sup>-1</sup>) | $G(t^*)/G(0)$ |
| --- | --- | --- | --- | --- | --- |
| A | 2427 | 330 | 433 | 1.14 | 2.11 |
| B | 3128 | 422 | 378 | 0.76 | 1.86 |
| C | 625 | 92 | 622 | 4.8 | 2.05 |

At this point we can investigate the influence of the parameters from Table 2 on the aggressiveness of the cancer. We select as the primary marker the time *t*\* that it takes for *B*(*t*) to reach 100 g. For example, increasing the glucose competitiveness of the cancer *α* results in shorter values of *t*\*, see Figure 4, left panel. Similarly, *t*^*^ is a decreasing function of the cancer glucose consumption rate *k_B_*, see Figure 4, right panel, and of the efficiency of glucose use by the cancer *c_B_* (simulations not shown). The same pattern holds when the parameters of a diabetic patient are varied (simulations not shown).

**Fig. 3.**
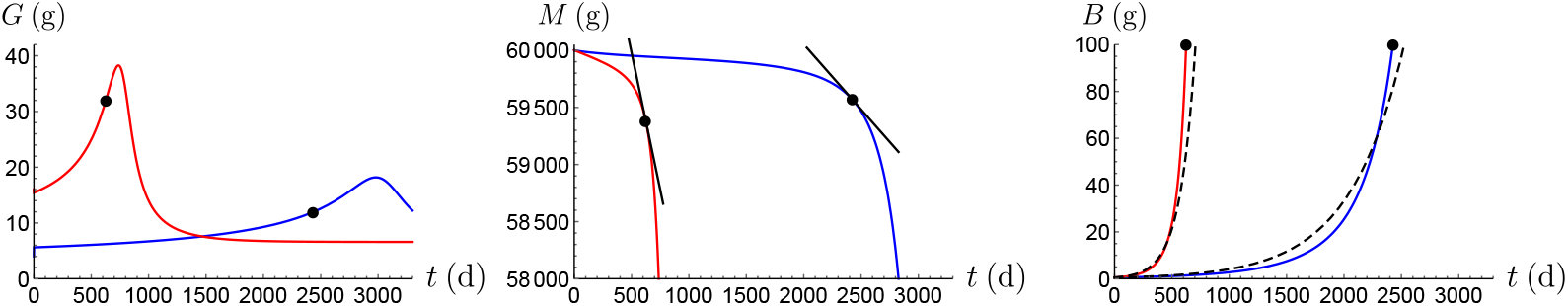
A plot of *G*, *M* and *B* for a non-diabetic patient (blue, column A of Table 1, scenario “A” again) and a diabetic patient (red, column B of Table 1, scenario “C”). For the diabetic patient we have *t*^*^ = 625 d. The cancer parameters are those listed in Table 2.

**Fig. 4.**
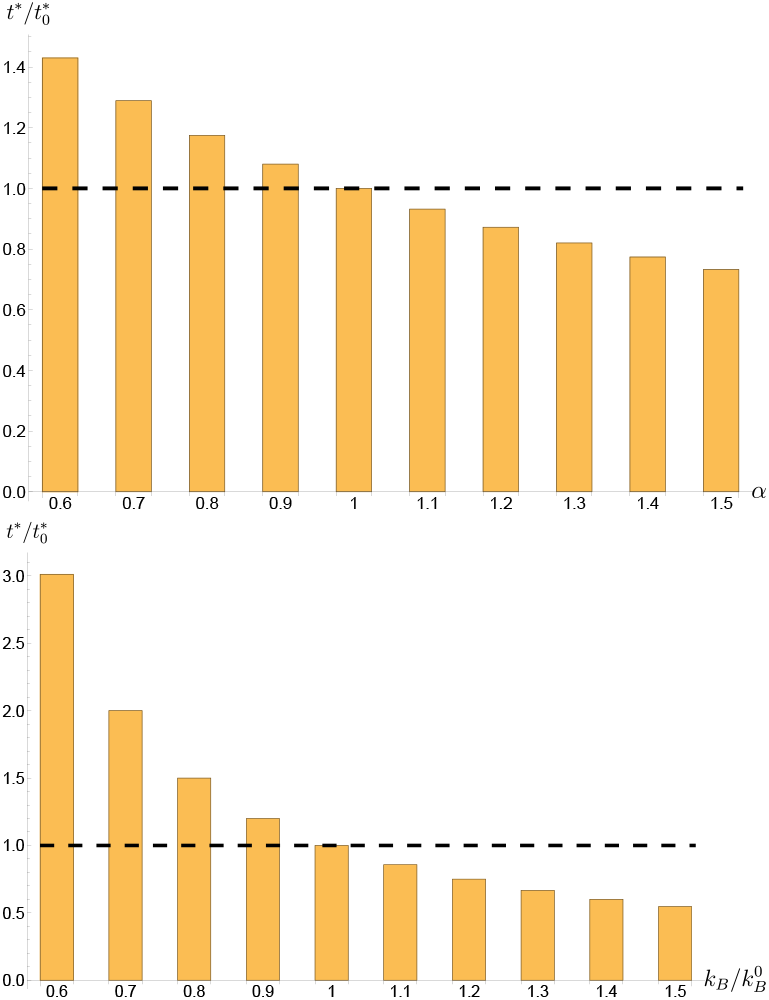
The relative eruption times *t*\* as a function of the cancer’s glucose competitiveness *α* (left; reference value is 1) and as a function of the cancer’s consumption rate *k_B_*. The cancer-unrelated parameters are those for a non-diabetic individual. The aggressiveness of the cancer increases from left to right.

Furthermore, our model allows to test strategies of using anti-diabetic drugs or dietary changes in supportive treatment of cancer patients. Three different approaches can be distinguished, namely

1. “type K” drugs that increase the uptake of glucose in healthy cells and hence in our model increase *k_M_*. These include most of the glucose-lowering medicines for type 2 diabetes known as biguanides, such as metformin that decreases glucose production in the liver and increases the body’s sensitivity to insulin. Metformin is often combined with other drugs for type 2 diabetes, such as glipizide, a sulfonylurea that increases release of body-produced insulin. In this case, both drugs function as “type K” drugs to cause a greater increase in *kM*. Supplemental insulin also increases *k_M_* and therefore belongs to this category.
2. “type M” drugs that inhibit the reabsorption of glucose in the kidneys, and hence in our model increase *m*. These include sodium-dependent glucose cotransporter 2 (SGLT2) inhibitors.
3. “type R” strategies that decrease the dietary glucose intake and in our model decrease *r*. Such a strategy can also include appetite-reducing drugs.

It needs to be recalled that changing any of these parameters has implications on the values of *k_M_* and *d_M_*, as these were determined using the equilibrium conditions (7) and (8), respectively. A change in *k_M_* in turn has further implications if the relationship (9) is assumed. With this in mind, our model provides guidance on what to expect from the different mitigation strategies and whether synergistic effects are possible. The simulations in Figure 5 suggest that a type *K* drug has a substantial effect in increasing *t*^*^ if relationship (9) is **not** in force. The effect of a type *K* drug is also more pronounced in a cancer patient without diabetes. In numbers, an increase of *k_M_* by a factor of 1.3 increases *t*\* by a factor of 1.62 in a non-diabetic patient but only by a factor of 1.36 in a diabetic patient. On the other hand, for a reduction of *r* to have an effect on *t*^*^ it is necessary for the relationship (9) **to be** in force, see Figure 6. Finally, an increase of *m* has a negligible effect on *t*^*^ (simulations not shown).

**Fig. 5.**
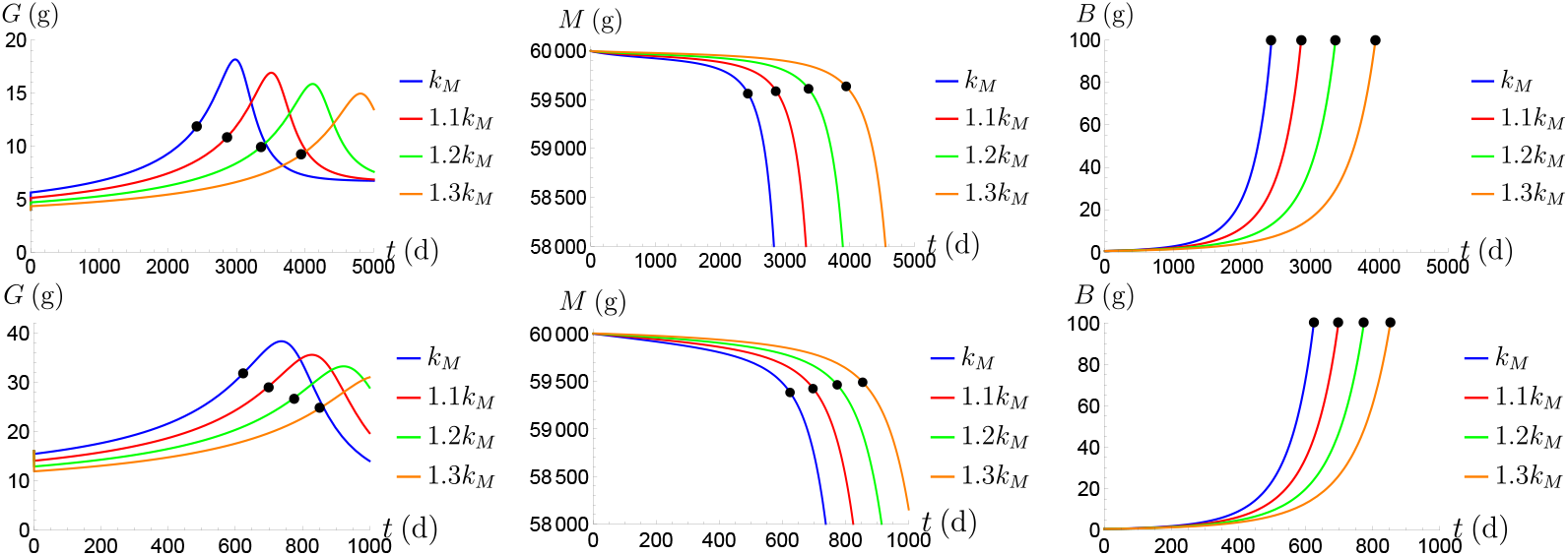
Plots of G, *M* and *B* for increasing values of *k_M_* when *k_B_* does not change in response. The cancer-unrelated parameters are those for a non-diabetic individual (top row) respectively for a diabetic individual (bottom row). The times *t*^*^ are marked by black dots.

**Fig. 6.**
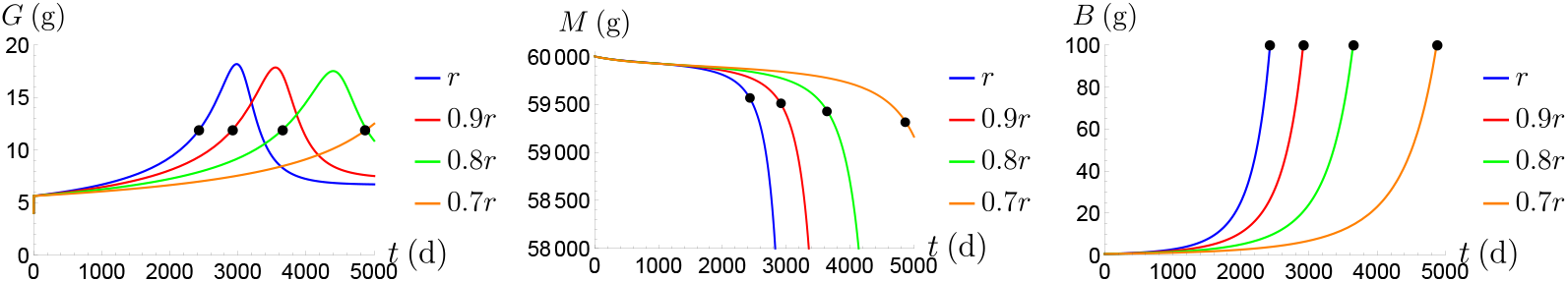
Plots of G, *M* and *B* for decreasing values of *r* when *k_B_* does change in response. The cancer-unrelated parameters are those for a non-diabetic individual. The times *t*^*^ are marked by black dots.

## 4 Discussion

In the following we gauge the model predictions from Figures 2 and 3 and Table 3 against observations with an emphasis on clinical or experimental studies. Naturally, there are large ranges of values reported in the literature, depending on the types of cancer and the goals of the particular studies. For example, in a clinical study [22] report doubling times of primary lung cancer between 30 and 1077 days, with a mean of 163 days. Similarly, the meta-study [23] lists a range of breast cancer doubling times of 150-250 days which has remained stable over the past 80 years. These ranges overlap with our values from Table 3 although the means are smaller than our values in non-diabetic cancer patients (scenarios A and B). Clearly, this depends on the choice of the cancer parameters in Table 2. Thus in future work these will need to be adapted to a particular type of cancer that is being studied. In recent work, [5] observed elevated fasting glucose levels in breast cancer patients of (mean value) 125.3 g/dL compared to 110.6 g/dL for the control group. This ratio of 1.13 is somewhat smaller than our values, but the average daily glucose levels could exhibit a greater difference between cancer and control groups due to the reported effect of cancer cells to suppress both basal and glucose-stimulated insulin secretion. Notably, all these parameters can be adjusted in future work to allow personalized modeling of tumor growth and response to interventions.

The time *t*^*^ can be interpreted as the time between inception of the cancer and its detection or the appearance of clinical symptoms. In our implementation, *t*\* depends on the choice of B(0) (here 0.5 g), and the threshold for detection (here 100 g). The time from emergence of a cancerous founder cell to clinical detection of the primary tumor is generally believed to be several years to a decade [24]. Such estimates are based on the fact that often metastases are present already at time of detection and these require a sequence of mutations. The time *t*^*^ is particularly valuable when comparing different scenarios. In our simulations we observe a reduction of *t*^*^ in a diabetic patient to about 25 % of its value in a non-diabetic patient (compare scenarios A and C). Diabetes or hyperglycemia are associated with a significantly higher risk of all-cause mortality in patients with NSCLC [25]. Similarly, the meta study by [26] indicates an increased cancer mortality in patients with an increased fasting glucose levels. There is a broad consensus that diabetes poses a general higher risk of developing cancer [27].

[28] report that in stage III and IV non-small cell lung cancer (NSCLC) patients weight loss occurs at a rate of 59 ±3 g/week. This is somewhat larger than what our model predicts, but it may also result from full scale cachexia, the involuntary loss of skeletal muscle and adipose tissue caused by profound metabolic rewiring of normal tissues during the late stage of the disease.

Therapeutic intervention is possible by re-purposing anti-diabetic drugs, even in cancer patients that had no earlier diagnosis of diabetes. Metformin decreases glucose production by the liver, increases insulin sensitivity of tissues, and further reduces appetite and caloric intake. Hence it is possible to view it as a type *K* drug as it was introduced above. In contrast, SGLT2 inhibitors (also known as gliflozins) such as empagliflozin increase the excretion of glucose through the kidneys. Whether type *K* drugs or the type R strategy have any impact on *t*^*^ depends on if and how the cancer glucose consumption rate *k_B_* changes as a response. If *k_M_* increases and *k_B_* stays the same, then *t*^*^ increases, see Figure 5. If *k_M_* decreases due to a decrease of *r* then *k_B_* needs to decrease as well, and then *t*^*^ will increase, see Figure 6. At present we are required to use equilibrium conditions such as (7) and (8) since we do not have access to all parameters (of the cancer-free model) from the literature. Whether the crucial connection (9) is solely a choice of our numerical implementation or whether there could be some biological background for it remains a question for future research.

Metformin has shown some improved survival rates in colorectal cancer, while use of insulin is associated with worse survival [9]. Metformin shows positive outcomes for pancreatic, colorectal, breast and lung cancers. On the other hand, there is no significant improvement in prostate, bladder, thyroid, renal, head and neck, esophageal, and hepatocellular cancers. [29] list various types of malignancies and whether they are associated with increased incidence of non-insulin dependent diabetes mellitus. We have simulated only an “abstract” cancer, but we have indicated how parameters can be adapted to match the characteristics of different types of malignancies. Beyond that, there is also the possibility of differences between patients with the same type of cancer. An elevated glucose consumption rate *k_B_* could be derived from a PET scan. It has been well-recognized that cancer cells in different tumors and even within the same tumor are heterogeneous, which is a combined result of the diverse tumor-intrinsic genetic and epigenetic factors (e.g., cancer cells’ abilities to consume glucose and suppress healthy tissues’ consumption) and host factors (e.g., status of glucose homeostasis). On the other hand, patients with similar tumor characteristics and host conditions could still have different prognoses. Therefore, those measurable tumor and host factors, together with other unknown or stochastic factors, together define the phenotypic characteristics and behaviors of cancer. In this regard, future modeling would benefit from tuning the general parameters determined herein using a large number of diverse tumors, and could include one or more terms reflecting a stochastic process.

## Conflict of interests/data sharing

The authors declare no conflict of interest. Data sharing is not applicable to this article as no datasets were generated or analysed during the current study.

## Acknowledgments

Jonathan Doria and Chinh Nguyen received awards from the Support for Undergraduate Research Fellows (SURF) program at the University of Wisconsin - Milwaukee. Noah Salentine was supported by the Ronald E. McNair Postbaccalaureate Achievement Program at the University of Wisconsin - Milwaukee. Shizhen Emily Wang was partially supported by National Institutes of Health grants R01CA218140 and R01CA266486. The funding bodies had no influence on study design, collection, analysis and interpretation of results. We thank Dr. Wei Ying (University of California San Diego) and two unknown readers for valuable comments.

